# Retinotopic maps of visual space in the human cerebellum

**DOI:** 10.1101/455170

**Authors:** D.M. van Es, W. van der Zwaag, T. Knapen

## Abstract

While the cerebellum is instrumental for motor control, it is not traditionally implicated in vision. Here, we report the existence of 5 ipsilateral visual field maps in the human cerebellum. These maps are located within the oculomotor vermis and cerebellar nodes of the dorsal attention and visual networks. These findings imply that the cerebellum is closely involved in visuospatial cognition, and that its contributions are anchored in sensory coordinates.

---

The purported role of the cerebellum has shifted from one that is exclusively sensorimotor-related to one that encompasses a wide range of cognitive and associative functions^1^. In fact, the majority of cerebellar cortex is functionally connected to a range of cognitive and associative cerebral networks^2^ and is coherently activated by cognitive and affective tasks^3,4^. Within sensorimotor areas of the cerebellum, a representation of the body (or ‘homunculus’) characterizes functional organization^5^. Yet, in the remaining cerebellar cognitive and associative networks, functional organization remains less well understood.

In cerebral cortex^6^^-^^8^ and subcortex^9^, orderly progressions of the visual field characterize functional organization in brain areas important for visual perception and cognition. Some of these areas are referred to as the dorsal attention network^10^, which was recently shown to have cerebellar counterparts^11^. One of these nodes was suggested to topographically encode visual field location^12^.

Here, we used the HCP retinotopy dataset^13^ to address the question of cerebellar visuospatial organization. The dataset contains population receptive field (pRF)^14,15^ parameters for the whole brain fitted on high-resolution 7T BOLD responses to visual retinotopic stimulation during fixation. The pRF model describes voxels’ visual field response preferences with a limited set of spatial parameters. The pRFs were fitted on data from individual subjects (N=181), and on an across-subject time course average (HCP ‘average subject’). As BOLD SNR is relatively low in the cerebellum this dataset provides an unprecedented opportunity to address cerebellar retinotopic organization, which we here use to identify 5 retinotopic maps in 3 separate cerebellar clusters. Fig. 1 shows two example pRFs in the cerebellum, as well as pRF polar angle in the cerebellar volume.

**Fig. 1:**
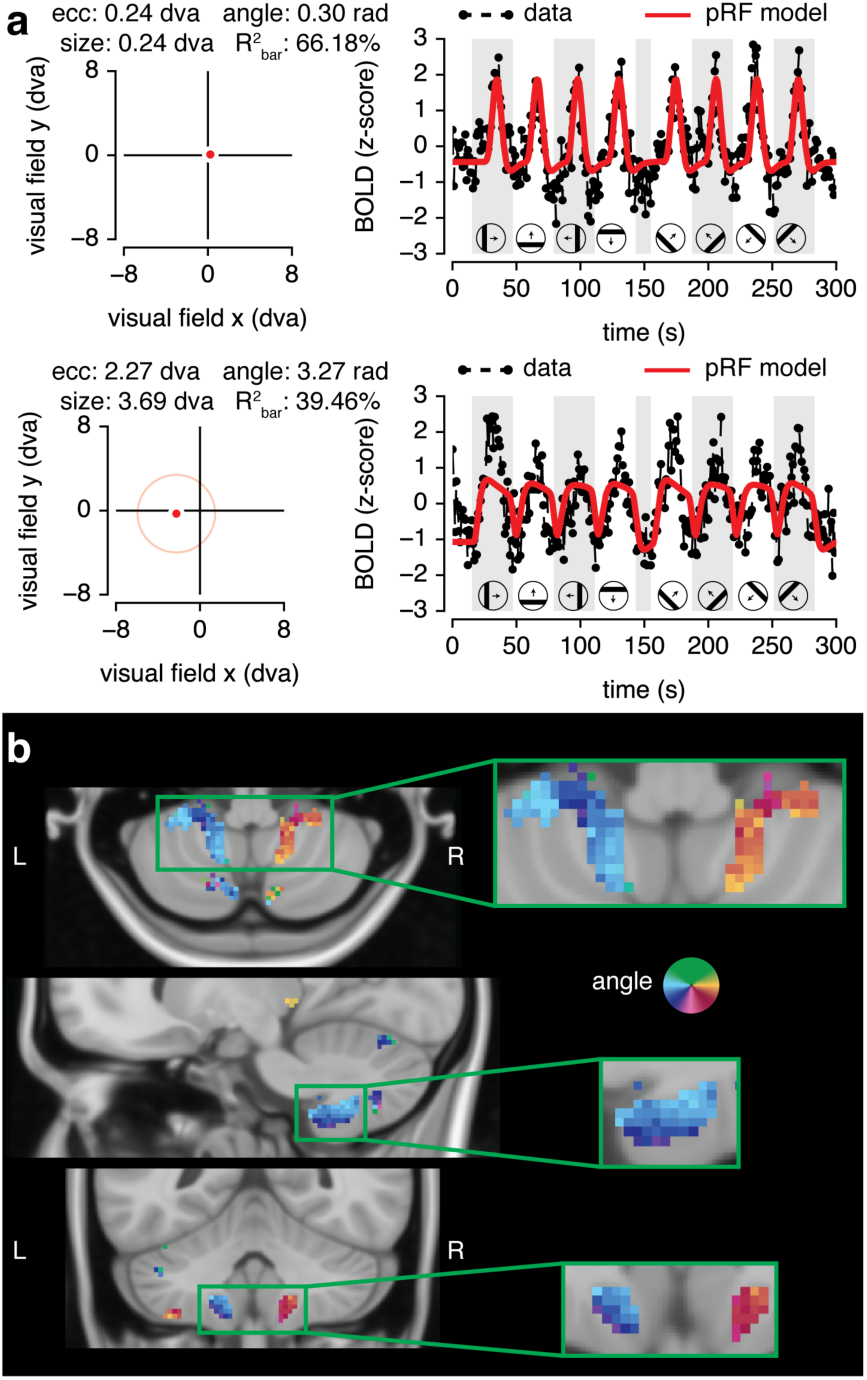
pRF fits in cerebellum. **a**, Example pRF profiles and fits for two voxels with different eccentricities and sizes. The dot in the visual field plot (left) indicates pRF center, the circle indicates pRF size at one standard deviation. The example time course (right) is the average across the two bar-stimulus runs. Explained variance displayed here (R^2^_bar_) is calculated across the fits and time courses shown. dva: degrees of visual angle. **b**, pRF polar angle in the volume. Insets highlight a cluster with ipsilateral progressions of the visual field (see Supplementary Fig 1 for all clusters). Data shown is from the HCP ‘average participant’.

To better inspect the topographic structure of visual-spatial representations in cerebellum, we projected pRF parameters for each cerebellar voxel onto a flattened representation of the cerebellum^16^ (Fig. 2a-d; see Supplementary Figs. 1 and 2 for results in the volume, Supplementary Fig. 3 for individual subject results, and Methods). This revealed three clusters where the pRF model explained considerable variance (see Supplementary Fig. 4 for the voxel selection procedure). We refer to the clusters as OMV, VIIb and VIIIb (see Fig. 2 g-h upper panels). Fig 2c shows that the distribution of pRF centers within each cluster is characterized by representations of the ipsilateral visual field. This is in contrast to the contralateral visual field representations in subcortical and cortical retinotopic areas^13^ (see Supplementary Fig. 2). Yet, it matches the ipsilaterality of the cerebellar homunculi^5^, resulting from midline crossing of cerebellar connective fibers in the pons^17^. Quantifications of the progressions of polar angle (Fig. 2g) reveal a double representation of the lower visual field in OMV and VIIIb, as is common in cerebral visual cortex. Finally, smooth variations in preferred eccentricity take place in the direction roughly orthogonal to the direction of polar angle phase reversals, again mirroring the organization of cerebral visual cortex (Fig 2h). Fig 2. e-f provides a visual model summary of these retinotopic properties.

**Fig. 2:**
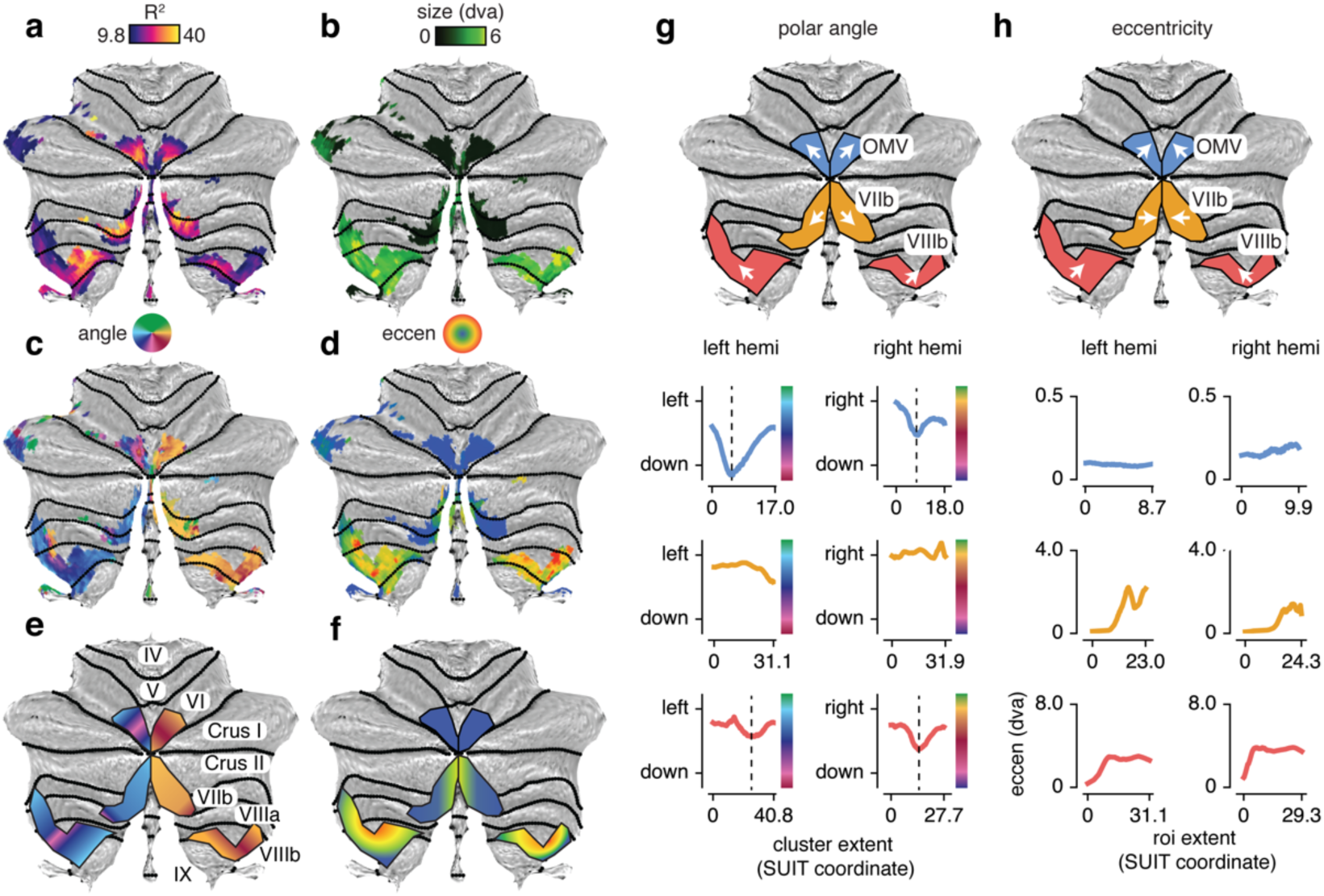
pRF parameters projected onto a flattened cerebellar representation. Flattened representation of pRF explained variance (**a**) size (**b**), polar angle (**c**), and eccentricity (**d**) reveal three retinotopic clusters in the cerebellum. Projections of pRF polar angle (**g**) and eccentricity (**h**) along the direction indicated by the white arrows reveal double representation of the lower visual field in OMV and VIIIb (dashed vertical lines demarcate polar angle reversals). Summarized representation of pRF polar angle (**e**) and eccentricity (**f**). Data shown is from the HCP ‘average participant’. See Supplementary Fig. 3 for individual subjects.

We next analyzed whether standard retinotopic properties (such as overrepresentation of the fovea and a strong correlation between pRF eccentricity and size^14^) were also present in the cerebellar visual field maps (Fig. 3). As Fig. 2g revealed double representations of the visual field in OMV and VIIIb, we split these clusters into a medial and lateral portion along their respective polar angle reversals (Fig 3a). Visualizing the eccentricity distributions (Fig 3b) quantifies the observation described above that eccentricity coverage is peri-foveal in OMV, extends somewhat into the periphery in VIIIb, and covers the full range of stimulated eccentricities in VIIIb. Importantly, Fig 3c reveals clear increases in pRF size with increasing eccentricity. Finally, Fig 3d highlights (1) the strong ipsilateral visual field representations, and (2) a strong overrepresentation of the lower visual hemifield in VIIIb. Together, this shows that the cerebellar visual field maps follow both known properties of retinotopic organization, albeit with unique idiosyncrasies (i.e. ipsilaterality, and strong overrepresentations of the fovea in OMV and of the lower visual field in OMV_lat_ and VIIb).

**Fig. 3:**
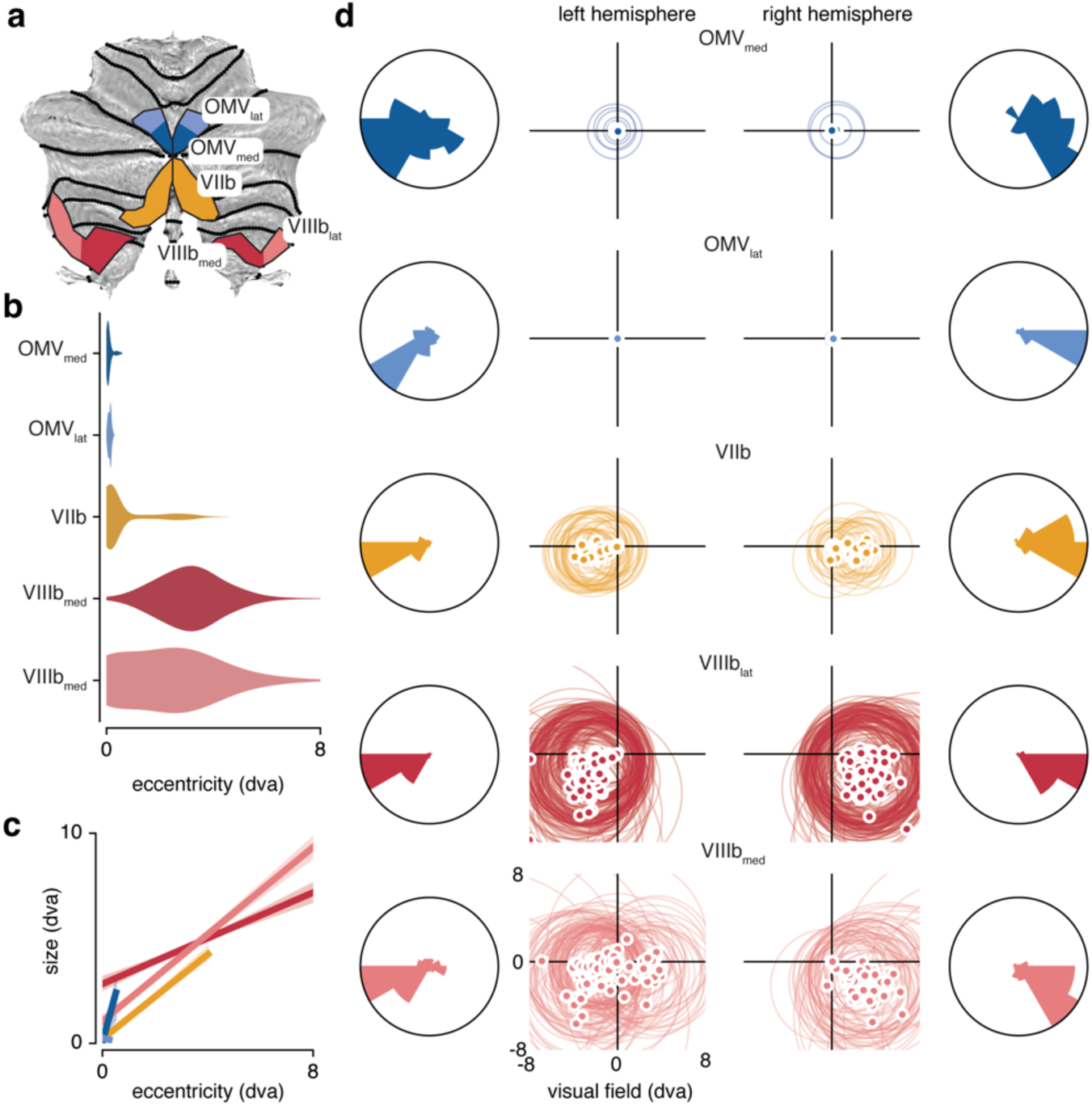
Distribution of pRF properties within cerebellar visual field maps. **a**, Legend. **b**, Distribution of pRF eccentricities. **c**, pRF eccentricity-size relations. Lines indicate linear regression fits with 95% confidence intervals across voxels as shaded regions. **d**, Distribution of pRFs throughout the visual field. Dots indicate pRF centers; circles indicate pRF size (one standard deviation). The polar histograms depict pRF center distributions. This shows (1) strong ipsilaterality in all maps, and (2) strong overrepresentations of the fovea in OMV and of the lower visual field in VIIIb. Data shown is from the HCP ‘average participant’. See Supplementary Fig 3. for individual subjects.

The oculomotor vermis (OMV) is implicated in the deployment of spatial attention and in the generation and adaptation of saccades^18^. Direction selectivity of OMV Purkinje cells was shown to arise as a function of saccade error direction, and are also organized along an anatomical gradient^19^. In addition, OMV neurons encode saccade amplitude by the duration of a population response rather than by amplitude tuning^20^. This could potentially explain the incomplete eccentricity coverages we find in the OMV cluster.

The anatomical location and extent of clusters VIIb and VIIIb overlap closely with cerebellar nodes of the dorsal attention network^2^. Visual field coverage was strongly biased to the lower visual field in VIIIb. Behavioral performance is known to be superior in the lower compared to the upper visual field for stimuli that are associated with visuomotor coordination^21^. In addition, activity in area VIIIb^22^ was shown to be related to (the observation of) reaching and grasping movements. This suggests that cluster VIIIb may be involved in the integration of visuospatial information for the guidance of effector movements.

Our results uncover 5 visual field maps in three retinotopically organized clusters in the cerebellum. This implies a much closer involvement of the cerebellum in visuospatial cognition than is classically assumed.

## Methods

### Data set

The pRF results presented in this manuscript are part of the 7T HCP retinotopy dataset. Please see the accompanying publication^13^ for details on data collection and model fitting procedures, and for links to the online repositories. Briefly, 181 subjects performed a discrimination task at the fixation mark while viewing expanding and contracting rings, rotating wedges and traversing bar stimuli filled with fast-changing, random visual stimuli, for a total of approximately 30 minutes scan-time. The maximum eccentricity for these stimuli was 8 degrees of visual angle. Visual selectivity for each ‘gray-ordinate’ is modeled as a population receptive field (pRF) model. This is a uniform Gaussian distribution with free parameters of center (x and y), size (standard deviation), amplitude (with fixed sub-additive normalization constant of 0.05) and a baseline parameter.

### Voxel selection procedure

In order to examine voxels that respond robustly to retinotopic stimuli, we first dismissed voxels where the pRF model explained little variance (see Supplementary Figure 4, first column; thresholds determined in original paper^13^ at 9.8% for the average and 2.2% for the individual subjects). Second, the non-linear spatial transformations that were employed to align data across subjects resulted in activity from ventral visual cortex to be smoothed into the cerebellar cortex. We were able to identify these voxels as these voxels were located between the cerebrum and cerebellum, and as these voxels were characterized by stark deviations in pRF parameter values (polar angle, eccentricity, size and explained variance; see areas indicated by white ovals in Supplementary Figure 4). The resulting mask left many voxels with extremely low eccentricity and size, without clear polar angle progressions across voxels. We hypothesized the following as a generative mechanism for these voxels’ results. As subjects performed a task on the fixation mark, this task became periodically more difficult when the retinotopic mapping stimulus passed behind the fixation mark. This means that responses of voxels sensitive to cognitive effort expended to maintain fixation (in a space-invariant manner) are in fact well captured by an extremely small and foveal pRF model. We therefore included an additional and conservative ‘fixation mark’ mask of voxels that had *both* eccentricity and size smaller than 0.15 degrees of visual angle (see Supplementary Figure 4, third column).

### Cerebellar flatmaps

We used the SUIT toolbox^16^ to project pRF results from three-dimensional volume space onto a flattened two-dimensional representation. Note that this flattened representation is compressed in the vertical dimension relative to a flattened representation that takes into account microscopic folding of individual cerebellum anatomy^23^.

### Subject ranking

In order to provide an estimate of the stability of the retinotopic maps in individual subjects (Supplementary Fig. 3), we ranked subjects based on the median explained variance across voxels within the three retinotopic clusters determined in the average subject. In creating visualizations of polar angles in these subjects, we masked voxels that fell outside the three retinotopic clusters as identified in the average subject and that were below the individual subject explained variance threshold of 2.2% (as determined in the HCP retinotopy paper^13^).

### Code availability

The analysis code for creating the figures presented in this manuscript will be published, upon acceptance under www.github.com/daanvanes/hcp_cerebellum_retinotopy.

## Supplementary Figures

**Supplementary Figure 1:**
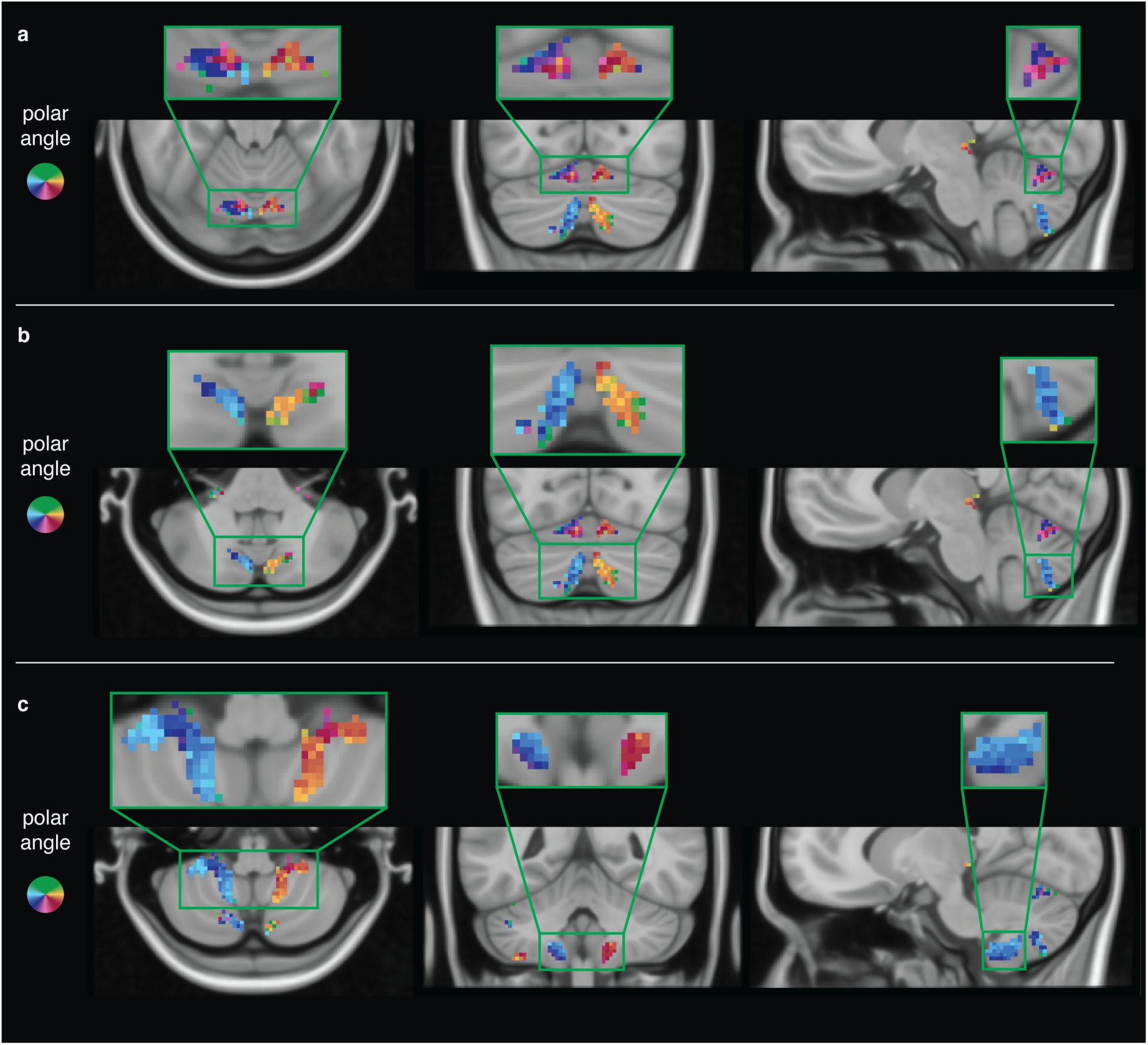
pRF polar angle in the three-dimensional volume representation. This figure depicts the three cerebellar clusters (Cluster VI (**a**), Cluster VIIb (**b**), and Cluster VIIIb (**c**)) in the volume using the 0.5mm MNI brain as anatomical reference. Insets are 200% magnification.

**Supplementary Figure 2:**
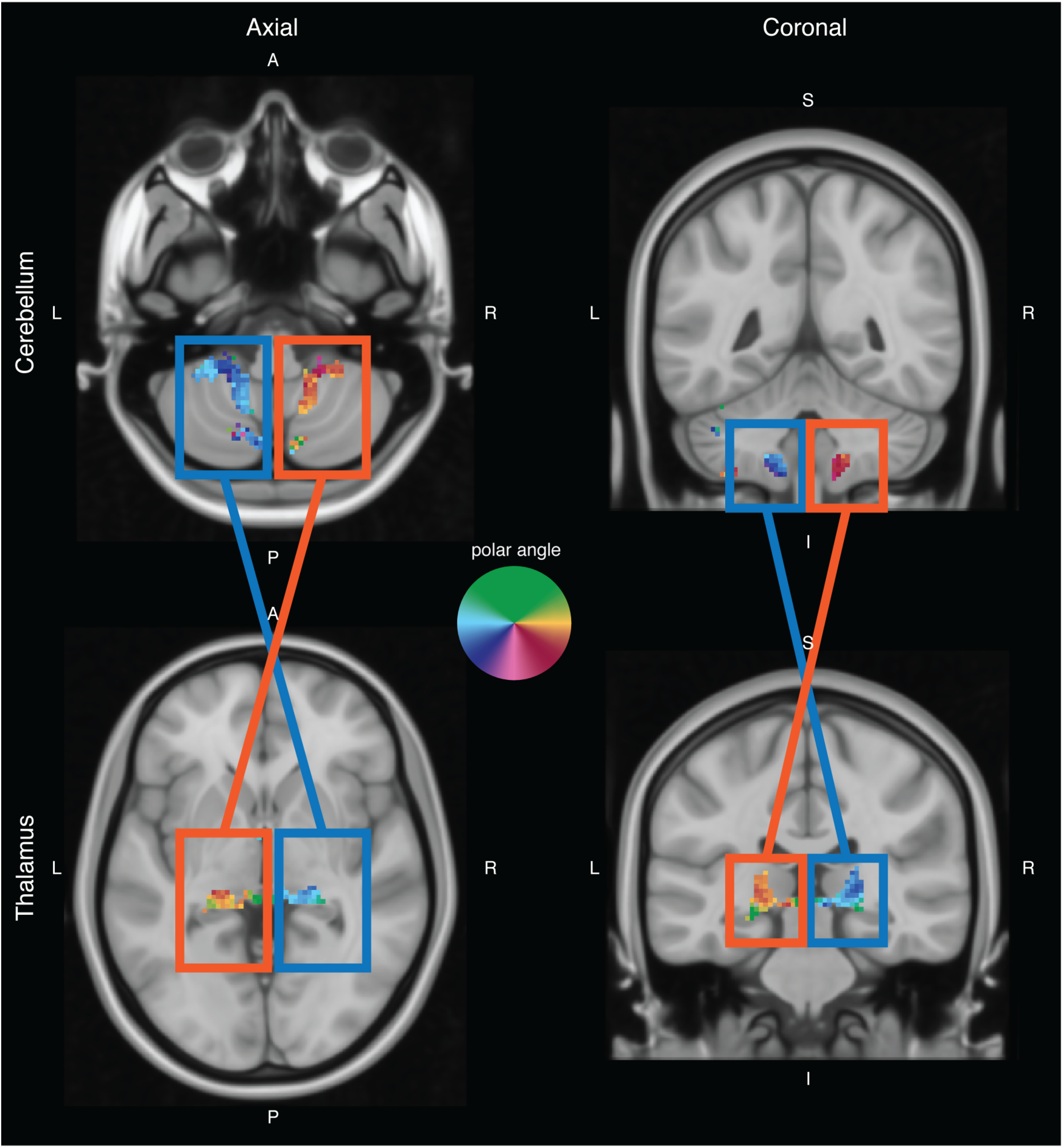
Polar angle representations in Thalamus and cerebellum. The ipsilateral vs. contralateral visual field representations in cerebellum vs. thalamus are evidenced by an inversion of blue and orange colors between left and right hemispheres.

**Supplementary Figure 3:**
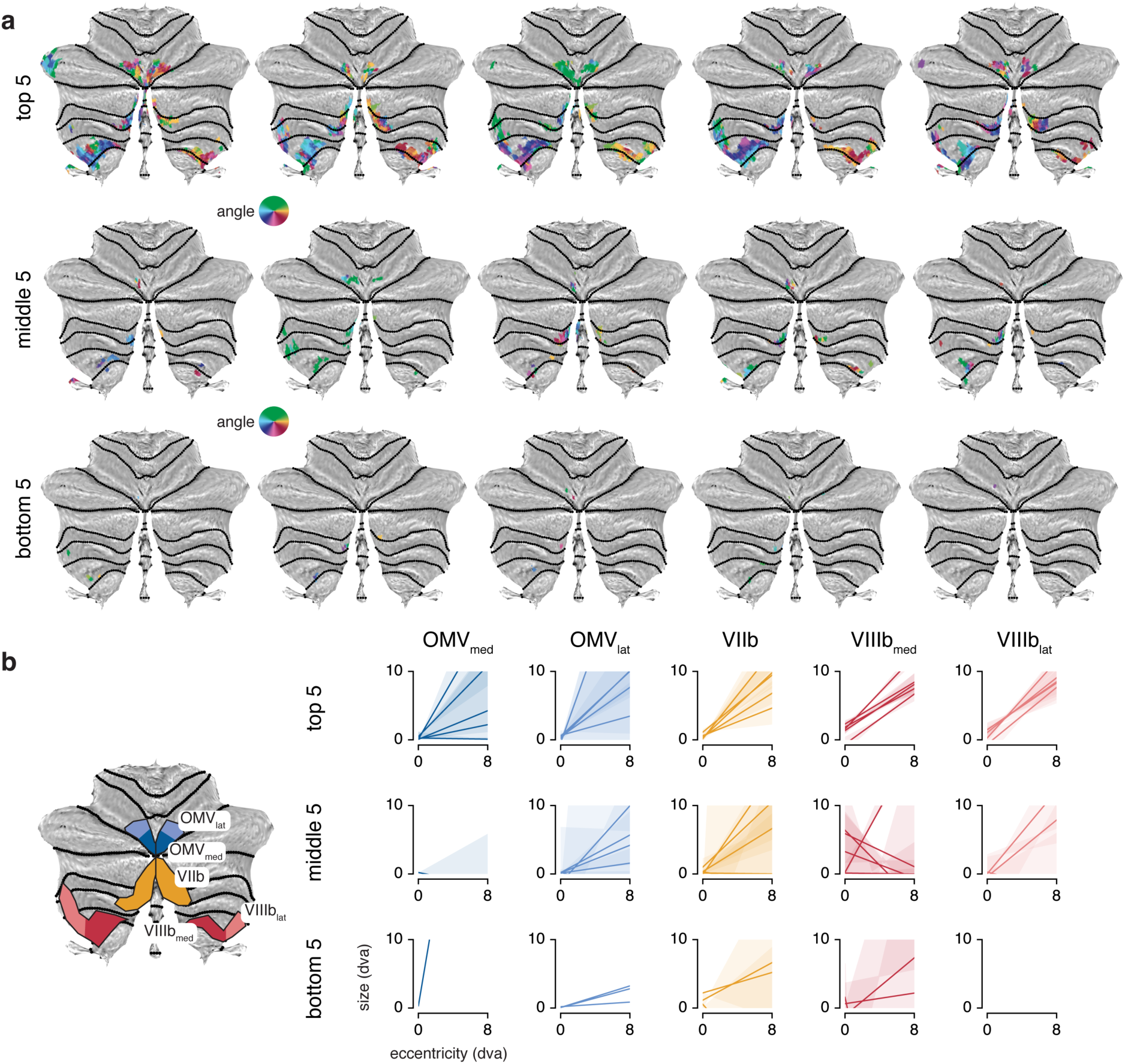
Individual subject results. **a**, Polar angle on the flattened cerebellar representation for the top, middle and bottom 5 subjects. For subject ranking see Methods. **b**, Eccentricity size relations for the top, middle and bottom 5 subjects. These results outline that retinotopic maps in cerebellar cortex are not readily identified in individual subjects with 30 minutes of retinotopic data at 7T. This is likely the result of the lower BOLD SNR in the cerebellum compared to the cerebrum1^5^, and highlights the unique opportunity to investigate these maps using the unprecedented statistical power of the HCP retinotopy dataset^13^. dva: degrees of visual angle

**Supplementary Figure 4:**
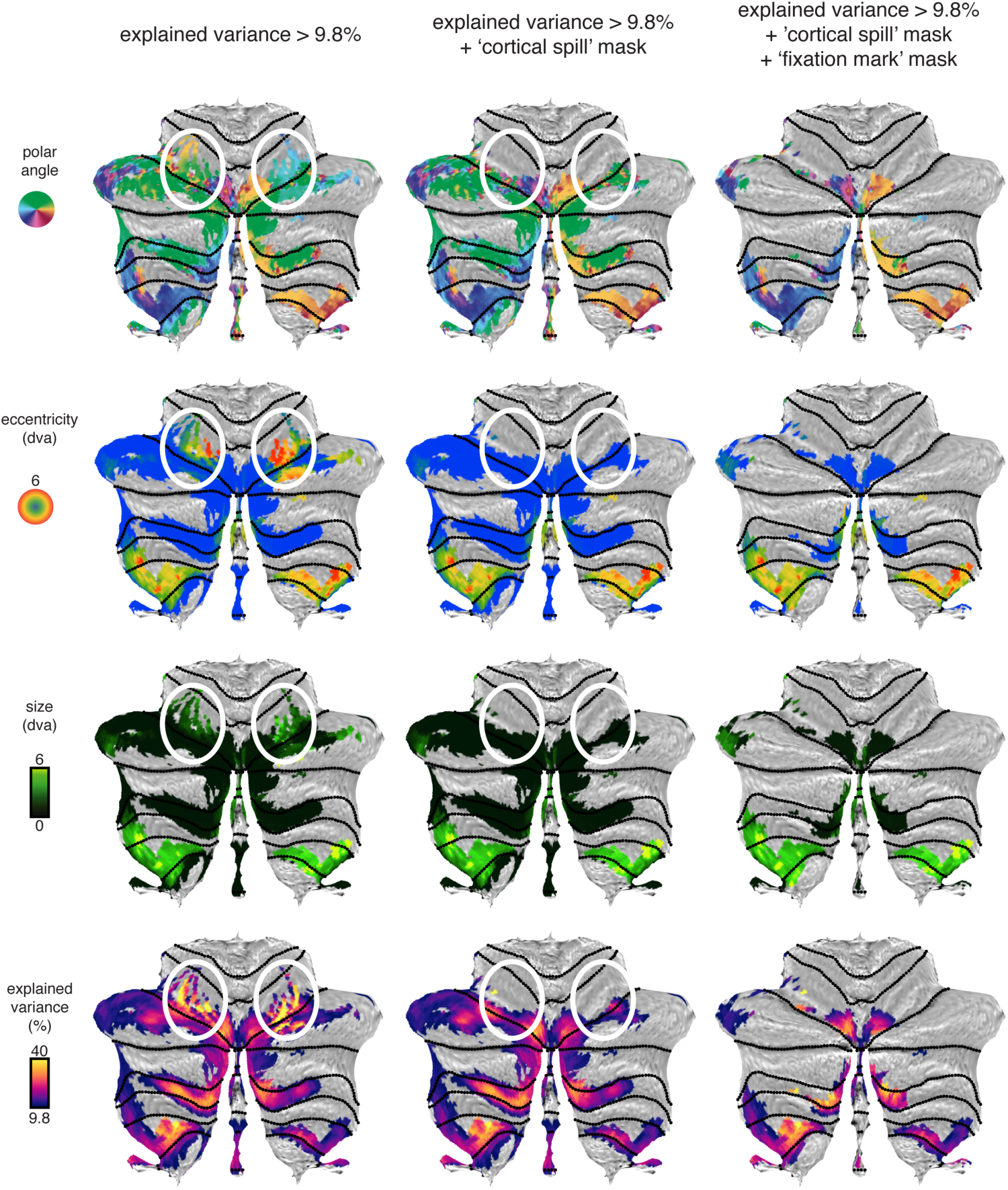
pRF polar angle, eccentricity, size and explained variance for different masks. pRF parameters projected onto flattened representations of the cerebellum for the different masking procedures (see Methods). The first row shows all voxels where explained variance is greater than the average participant threshold of 9.8% determined in the original paper^13^. The white ovals indicate voxels that are removed when deselecting voxels on the border between the cerebellum and cerebrum (the ‘cortical spill’ mask, second column). The third column shows voxels included in the final analysis. Here, voxels with a size and eccentricity smaller than 0.15 degrees of visual angles are additionally deselected (i.e. ‘fixation mask’). dva: degrees of visual angle.

